# The acute toxicity of Oxydemeton-methyl in zebrafish

**DOI:** 10.1101/119982

**Authors:** Guofeng Jia, Xiecheng Liu

**Affiliations:** Department of Health Management, Qingdao Municipal Hospital, Qingdao 266021, Shandong, China.

**Author notes:** Corresponding author: Dr. Guofeng Jia.

**Keywords:** Oxydemeton-methyl, toxicity, cytochrome P450, ROS, DNA damage, oxidative stress, zebrafish

## Abstract

Oxydemeton-methyl, is an organothiophosphate insecticide, which is widely used in agricultural and urban pest controls. It exists in the environment and a large amount bioaccumulation in the wildlife due to its strong water solubility and mobility. Although its potentially harmful effect on animals and humans, few studies have focused on the oxydemeton-methyl pollution in the environment. Zebrafish have been used for many years to valuate the pollution status of water and toxicity of chemicals. In the present study, we aimed to investigate the effect of oxydemeton-methyl on the expression level of liver microsomal cytochrome P450, on the activity of NADPH-P450 reductase and reactive oxygen species (ROS) generation in zebrafish. Adult male and female zebrafish were treated with different concentration of oxydemeton-methyl (10, 50, 100 μM) for 5, 10, 20 and 30 days. We found that the oxydemeton-methyl exposure significantly increased the P450 levels and the activity of NAPDH-P450 reductase. ROS generation and the DNA damage were augmented in a dose-dependent manner in the zebrafish. These results indicated that oxydemeton-methyl is able to induce strong oxidative stress and hence highly toxic to the zebrafish.

## Introduction

Oxydemeton-methyl is a widely organothiophosphate insecticide, which has primarily used to control pest [1–4]. Many countries have realized the high toxicity of oxydemeton-methyl to wildlife and made restriction or ban of the usage of this pesticide [5–7]. However, oxydemeton-methyl remains to be one of the most frequently used organophosphorus insecticides. The high water solubility and mobility led to significant harmful residues in the environment.

Studies on toxicity testing are usually conducted in mammalian models, such as rodents and rabbits [8, 9]. Whatever these tests are commonly expensive and require a large amount of animals. Fish, like many other animal species in the aquatic environment, has been widely used to investigate the toxicology of organophosphorus insecticides [10–13]. Zebrafish has been shown to be a valuable animal model to assess the effect of pollutants to the aquatic ecosystems [14–18]. Zebrafish is a useful experimental model for investigating vertebrate development because of its transparent embryos, low maintaining cost, conservation of key genes and signaling and easily genetic modification [19–25]. Hence, zebrafish has become increasingly common in compounds screening and drug discovery for evaluation of the toxicity mechanisms and also in drug selection and optimization [26–29]. The potential impacts of oxydemeton-methyl to the fish and the increased bioaccumulation effects of the toxicant in wildlife still need be further explored.

One of the most important manifestations in fish is oxidative stress. Increased ROS levels, antioxidant defense systems impairment and loss function of the oxidative self-repair can result in potential damage to fish [30–35]. The level of cytochrome P450 enzymes has been well established in fish as a monitoring indicator for evaluating of environmental contamination and ecotoxicology experiments [36–40].

In the present study, we examined the sensitivity of various biomarkers in zebrafish exposed to oxydemeton-methyl, thus gain a further understanding of the impacts of oxydemeton-methyl to the aquatic ecosystem.

## Materials and methods

### Chemicals

All chemicals (analytical standard) used in this study were purchased from Sigma-aldrich.

### Zebrafish husbandry

The wild type zebrafish (*Danio rerio*) were maintained at 28.5 °C on a 10-hours dark and 14-hours light cycle. All procedures were approved from the Qingdao Municipal Hospital Institutional Animal Care and Use Committee (2017N000105).

### Oxydemeton-methyl exposure

Six-month old adult male and female zebrafish were separated and housed in fish tanks. The body weight of male and female zebrafish were 0.45 ± 0.05 g and 0. 52 ± 0.08 g, respectively; and the length of male and female zebrafish were similar to 3.5 ± 0.3 cm. Both groups were treated with or without oxydemeton-methyl (10, 50, 100 μM) and samples were collected at 5, 10, 20 and 30 days post exposure. The fish were fed twice daily with commercial fish food and starved overnight prior to examination to avoid the effects of feces during the procedures of the assays. Half of the fish water was changed daily during the period of exposure to maintain the stable concentrations of oxydemeton-methyl.

### Protein measurement

The protein concentration was measured by using the Piece BCA Protein Assay Kit (Thermo Fisher Scientific) according to the manufactures’ instruction.

### Isolation of liver microsomes

The zebrafish were anesthetized in 0.4% tricaine (MS-222) and transferred to a moist sponge for surgery on ice. The livers were dissected and rinsed with ice-cold 1× PBS (137 mM NaCl, 2.7 mM KCl, 10 mM Na_2_HPO_4_, 1.8 mM KH_2_PO_4_, PH 7.4). The livers were transferred to ice-cold homogenization buffer (0.1 M sodium phosphate buffer, 1 mM EDTA, 0.1 mM DTT and 0.1 mM PMSF, PH 7.5) with 10% (v/v) glycerol and homogenized. The homogenate was centrifuged at 16,000 g for 20 min at 4 °C and the supernatant was centrifuged at 100,000 g for 1 hour at 4 °C. The microsomal pellet was collected in homogenization buffer with 20 % (v/v) glycerol and stored at -80 °C.

### P450 enzyme activities

The cytochrome P450 content and NADPH-P450 reductase activity were determined as described somewhere else [41, 42].

### Antioxidant enzymes activities

The homogenate was centrifuged at 16,000 g for 20 min at 4 °C, and the supernatant was collected and used for determination of the enzyme activity and protein concentrations.

By determining the inhibition of SOD to the photochemical reduction of nitroblue tetrazolium chloride (NBT), the SOD and CAT activity was measured as described previously [43–46].

### ROS production

The ROS production was determined according to the method described previously [47–49].

### Statistical analysis

Statistical analysis was performed using SPSS (IBM, Armonk, NY). Graphs were plotted using Prism Graph Pad software (6.0). Two-way ANOVA for repeated measurements used to determine the differences between duration and concentrations. All the values were presented as mean ± SEM. *P* ≤ 0.05 was considered statistically significant by Student’s t tests.

## Results

### Exposure to oxydemeton-methyl impaired cytochrome P450 and NADPH-P450 reductase activities

Compared to male fish treated with oxydemeton-methyl, the cytochrome P450 contents were higher in the female fish at the same exposed concentrations (Fig. 1). The P450 contents were increased and reached a maximum at the 10 days and then the induction showed a decreasing trend (Fig. 1). The P450 contents were higher at all experiment duration in the exposed fish compared to the controls.

**Figure 1.**
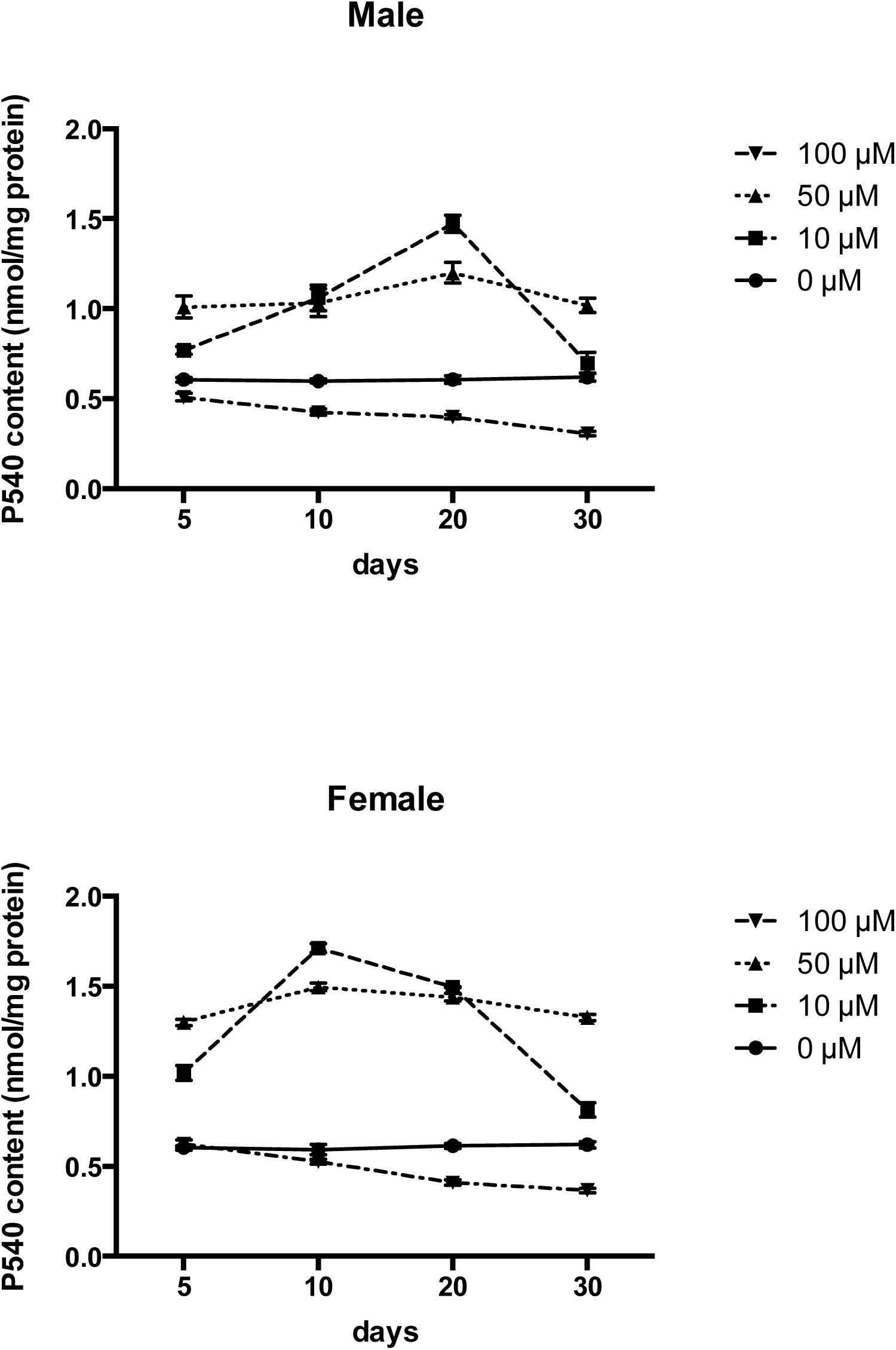
Effect of oxydemeton-methyl on the cytochrome P450 contents in zebrafish.

The NAPDH-P450 reductase (NCR) activity of oxydemeton-methyl treated fish was higher than the controls at all concentration during the exposure (Fig. 2). Oxydemeton-methyl significantly induced the NCR activity, and which was higher in the female than the male fish treated with the same concentrations. The NCR activity reached a maximum at 20 days both in the male and female fish exposed to oxydemeton-methyl (Fig. 2).

**Figure 2.**
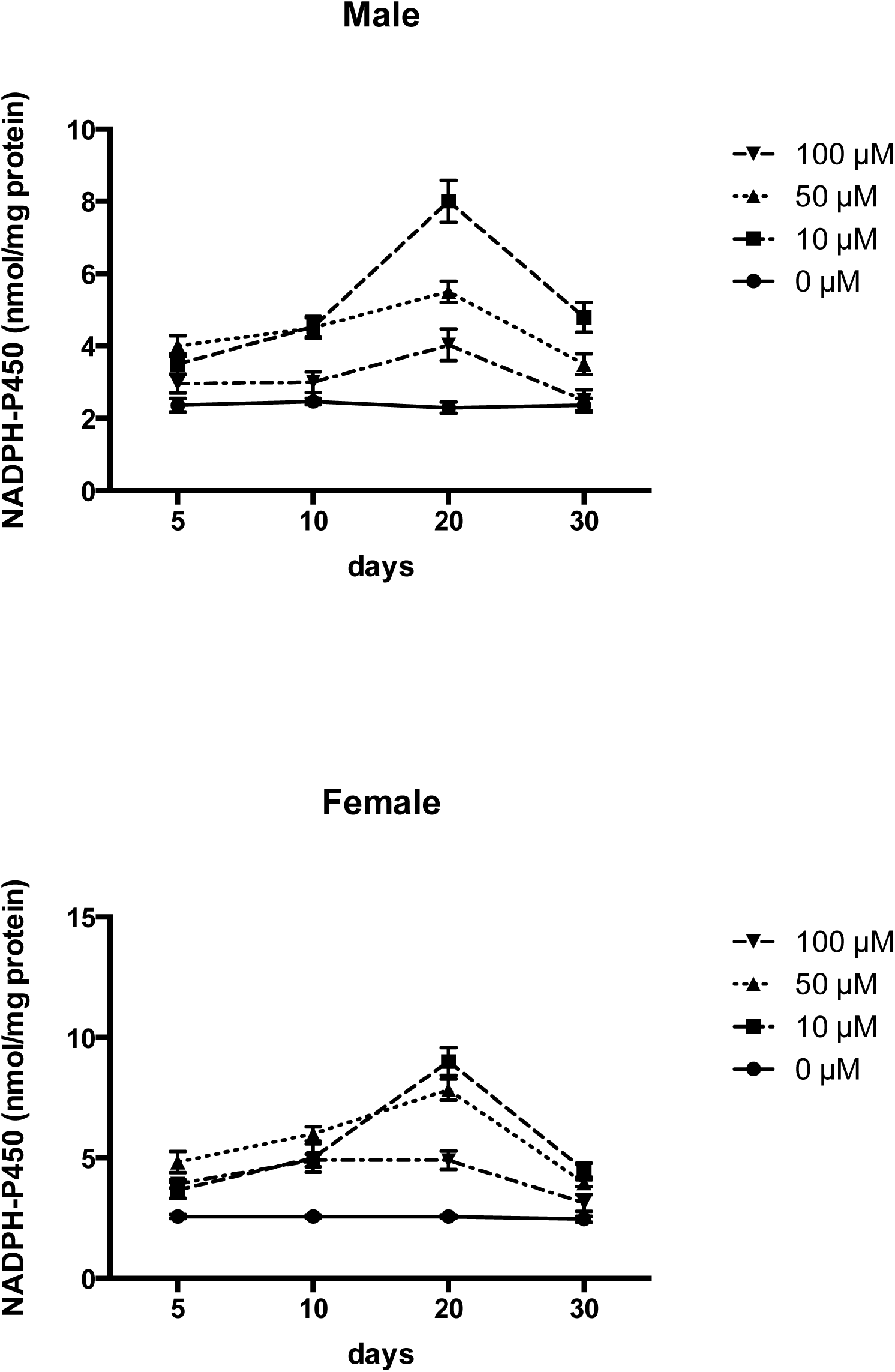
Effect of oxydemeton-methyl on the NADPH-P450 reductase activity in zebrafish.

### Exposure to oxydemeton-methyl inhibited anti-oxidative enzymes activity

During exposure, oxydemeton-methyl significantly inhibited the SOD activity in both male and female fish consistently over time (Fig. 3). The SOD activity of the female fish was higher then the same concentration treated male fish.

**Figure 3.**
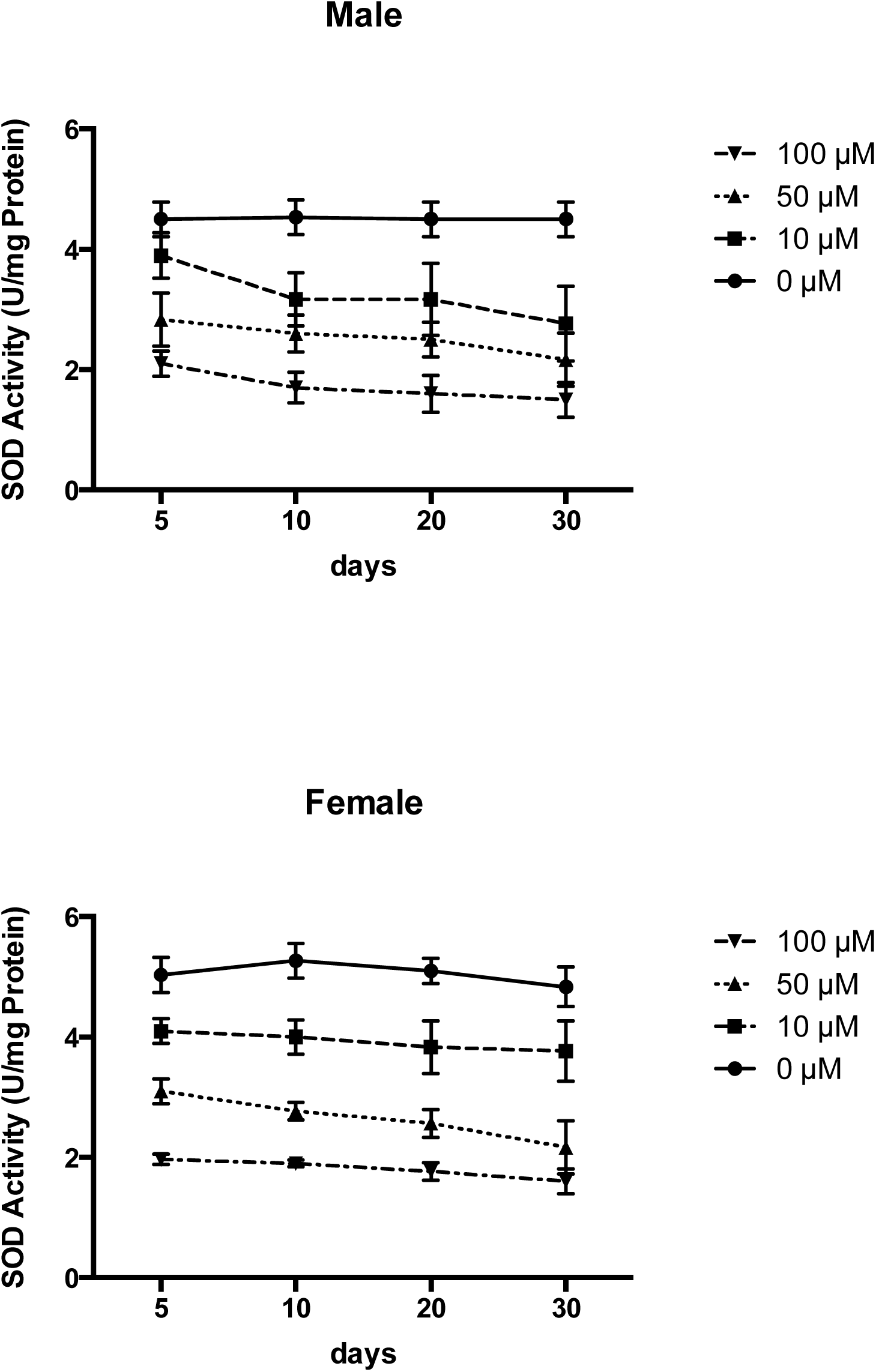
Effect of oxydemeton-methyl on the SOD activity in zebrafish.

The CAT activity was induced by low concentration of oxydemeton-methyl (10 μM) and inhibited by higher concentration and showed a consistent decrease during the exposure (Fig. 4). The CAT activity in the female fish was higher than in the male fish when treated with the same concentrations of oxydemeton-methyl, and which reached a maximum at 10 days. The CAT activity in the male fish reached the maximum at 20 days.

**Figure 4.**
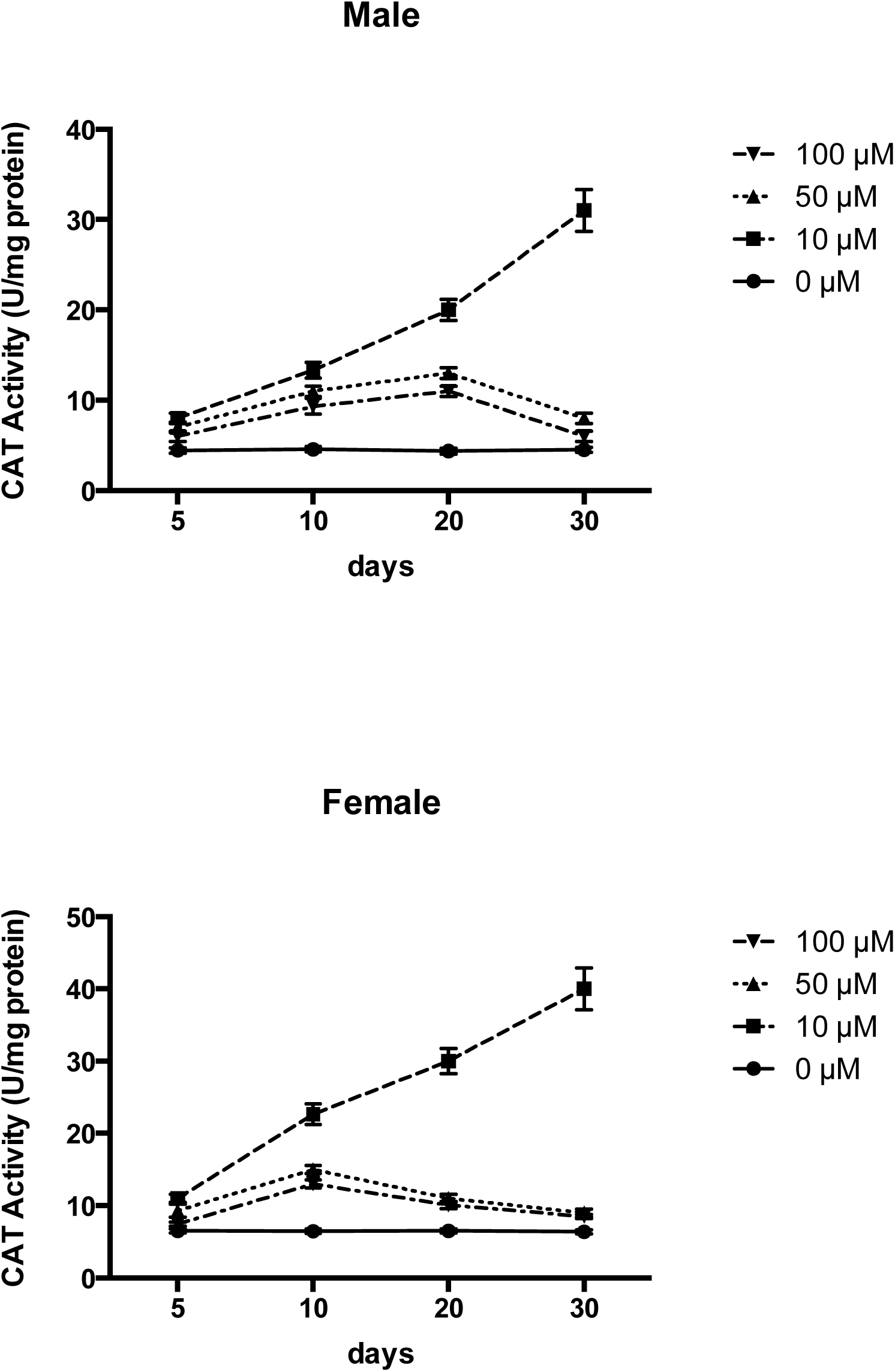
Effect of oxydemeton-methyl on the CAT activity in zebrafish.

### Effect of oxydemeton-methyl treatment on ROS production

The ROS production was significantly activated by oxydemeton-methyl treatment. Compared to the controls, the ROS levels in the exposed fish were higher, and had a tendency to increase at all oxydemeton-methyl concentration (Fig. 5). The ROS levels were higher in the female than the male fish, which may reflected some sex differences.

**Figure 5.**
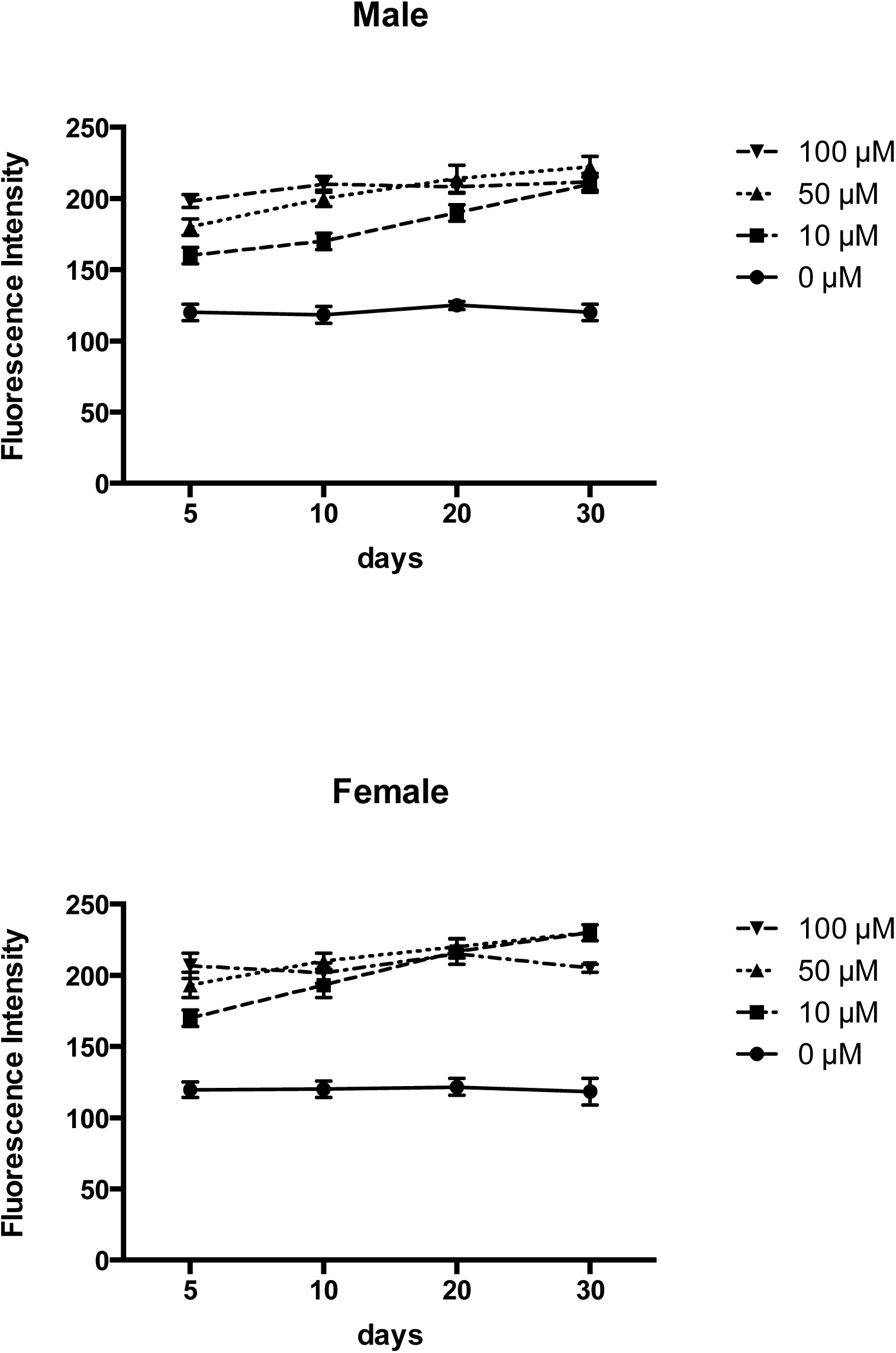
Effect of oxydemeton-methyl on the ROS production in zebrafish.

## Discussion

Our data showed that oxydemeton-methyl exposure could affect the cytochrome P450 and ROS contents, SOD NCR and CAT activities in the zebrafish. Cytochrome P450 enzymes activities are useful marks that can be used in environmental contamination biomonitoring and xenobiotic metabolism tests [50–52]. The P450 contents were impacted by oxydemeton-methyl treatment and the stimulation trend appeared an initial increase followed by a significant decrease over time. The alteration of NCR would affect the function of the monooxygenase system [53]. We found that cytochrome P450 affect the activity of NCR, which might be the role of P450 enzyme activity in detoxification.

The activity of SOD and CAT was significantly induced by oxydemeton-methyl treatment, which might due to oxydemeton-methyl increased ROX production in the exposed zebrafish. Increased CAT and SOD activities eliminate the redundant ROS and maintain ROS levels at a steady-state concentration [54, 55]. ROS, including a large amount of reactive chemically molecules derived from oxygen, are generated during normal physiological process in all aerobic organisms [56, 57]. ROS can directly damage cellular components and affect cell function [58–60]. Oxydemeton-methyl treatment significantly increased ROS levels in zebrafish, suggested its high toxicity which caused the organisms were not able to eliminate the exceed ROS. The generation of ROS might damage membrane lipids, DNA, protein metabolism and barbohydrate activities.

Our study showed that the cytochrome P450 and ROS content, SOD, CAT and NCR activities were higher in female than in male fish, that suggested the sex differences can affect enzyme activity. These results indicated that female zebrafish could be a good biological indicator in pollution evaluation. Our study could provide a basic theory to further studies of toxicity mechanisms of oxydemeton-methyl in animals.

